# Enhanced pre-frontal functional-structural networks to support behavioural deficits after traumatic brain injury

**DOI:** 10.1101/057588

**Authors:** Ibai Diez, David Drijkoningen, Sebastiano Stramaglia, Paolo Bonifazi, Daniele Marinazzo, Jolien Gooijers, Stephan P. Swinnen, Jesus M. Cortes

## Abstract

Traumatic brain injury (TBI) affects its structural connectivity, triggering the re-organization of structural-functional circuits in a manner that remains poorly understood. We studied the re-organization of brain networks after TBI, taking advantage of a computational method based on magnetic resonance imaging (MRI) including diffusion-weighted imaging in combination with functional resting state data obtained from the blood-oxygenation-level-dependent T2^*^ signal. We enrolled young participants who had suffered moderate to severe TBI (N=14, age 13.14 ± 3.25 years), comparing them to young typically developing control participants (N=27, age 15.04 ± 2.26 years). We found increased functional and structural connectivity within a cortico-subcortical network in TBI patient’s brains that involved prefrontal cortex, anterior cingulate gyrus, orbital gyrus and caudate nucleus. In comparison to control participants, TBI patients increased functional connectivity from prefrontal regions towards two different networks: 1) a subcortical network including part of the motor network, basal ganglia, cerebellum, hippocampus, amygdala, posterior cingulum and precuneus; and 2) a task-positive network, involving regions of the dorsal attention system together with the dorsolateral and ventrolateral prefrontal regions. We also found the increased prefrontal activation in TBI patients was correlated with some behavioural indices, such as the amount of body sway, whereby patients with worse balance activated frontal regions more strongly. The enhanced prefrontal activation found in TBI patients may provide the structural scaffold for stronger cognitive control of certain behavioural functions, consistent with the observation that various motor tasks are performed less automatically following TBI and that more cognitive control is associated with such actions.

## Abbreviations

COG: centre of gravity
COP: centre of pressure
DC: Directional control
DWI: diffusion weighted imaging
FACT: Fiber Assignment by Continuous Tracking
fMRI: functional Magnetic Resonance Imaging
FN: functional networks
ICA: independent component analysis
iPL: inverse path length
LOST: The Limits Of Stability Test
MRCs: most representative clusters
PPA: point process analysis
ROIs: regions of interest
RWST: Rhythmic Weight Shift Test
SC: structural connectivity
SN: structural networks
TBI: Traumatic brain injury

## 1. Introduction

Traumatic brain injury (TBI) disrupts functional and structural large-scale brain networks (the latter meant here as the white matter tracts connecting different brain regions), generally produced by a considerable impact on the head that damages to specific white matter tracts. Such lesions may provoke short or longer episodes of unconsciousness and may give rise to various behavioural deficits such as cognitive impairments, motor problems, emotional sequelae, etc. even years post-injury (Smith and Meany 2000; Caeyenberghs *et al.*, 2012; Ham *et al.*, 2012).

Over the past decades imaging techniques such as diffusion weighted imaging (DWI) and functional Magnetic Resonance Imaging (fMRI) have progressed our understanding of the physiopathology of TBI. In particular, recent advances in MRI techniques allowed analysing the injured brain and its correlation with behaviour from a network perspective, i.e., exploring the structural and functional connectivity of neuronal networks in vivo injury (Bonnelle *et al.*, 2011; Caeyenberghs *et al.*, 2012, Caeyenberghs *et al.* 2013, Caeyenberghs *et al.* 2014a; Ham *et al.*, 2012; Maki-Marttunen*et al.*, 2013; Sharp *et al.*, 2014, Sharp *et al.* 2015; Barbey *et al.*, 2015; Fagerholm *et al.*, 2015). Thus, for instance, diffusion weighted results revealed reduced structural connectivity and reduced network efficiency in TBI in relation to poorer cognitive functioning (Bonnelle *et al.*, 2011; Caeyenberghs *et al.*, 2014a; Kim *et al.*, 2014; Fagerholm *et al.*, 2015), and poorer balance control (Caeyenberghs *et al.*, 2012). Moreover, with respect to resting state functional connectivity, i.e., looking at regional BOLD interactions when the brain is at rest, multiple studies have reported TBI-induced alterations (Bonnelle *et al.*, 2011, Bonnelle *et al.* 2012; Hillary *et al.*, 2011; Sharp *et al.*, 2011, Sharp *et al.* 2014; Tarapore *et al.*, 2013), even in cases of mild TBI (Mayer *et al.*, 2011; Zhou *et al.*, 2012).

It has been also reported increased connectivity in TBI patients in relation to healthy participants within the default mode network (the most basal network at rest which plays an integrative role between other networks) (Sharp *et al.*, 2011; Hillary *et al.*, 2011; Palacios *et al.*, 2013), possibly acting as a compensatory mechanism for loss of structural connections (axonal injury). Importantly, TBI-induced changes in resting state functional connectivity appear to be able to predict, for example, the development of attentional impairments [4]. Additionally, Caeyenberghs and colleagues (Caeyenberghs *et al.*, 2013) revealed that combining information from structural and functional networks resulted in a better prediction of task-switching performance in TBI.

Although there is now sufficient evidence that TBI damages large-scale and emerging properties of brain structural and functional networks, and that the degree of network impairment is correlated with behavioural and cognitive deficits, the precise pattern of structural-functional circuit re-organization after TBI is still poorly characterized. Based on previous studies exploring how the brain reorganizes after TBI (Vakhtin *et al.*, 2013; Caeyenberghs *et al.*, 2014b; Drijkoningen *et al.*, 2015a), we have used a novel brain atlas (Diez *et al.*, 2015) to probe the working hypothesis that when structural networks are damaged and reorganized as a result of TBI, there is an associated reorganization of the corresponding functional networks, and vice versa. This hypothesis is in accordance with previous work emphasizing the strong mutual relationship between brain structure and function (Damoiseaux *et al.*, 2009; Park *et al.*, 2013; Diez *et al.*, 2015). As such, the proposed atlas is rooted in a common structure-function skeleton (Diez *et al.*, 2015), such that voxels belonging to the same region in the atlas have homologous functional dynamics and at the same time, they are structurally wired together. This has led to the discovery that both classes of networks (structural and functional) reorganize and enhance their connectivity in the prefrontal part of the brain, a region studied intensely in TBI patients, and associated with various behavioural deficits, including those pertaining to cognitive control and executive function (Levin and Kraus, 1994; Miller, 2000; Godefroy, 2003; Bonnelle *et al.*, 2011; Sharp *et al.*, 2011; Caeyenberghs *et al.*, 2014a).

## 2. Materials and Methods

### 2.1. Participants

A total of 41 young subjects were included in this study, 14 of whom suffered moderate to severe TBI (age 13.14 ± 3.25 years; 6 males and 8 females) and 27 age-matched control subjects who developed normally (age 15.04 ± 2.26 years; 12 males, 15 females). The TBI patients’ mean age at the time of injury was 10 ± 2.26 years, and the average time-interval between injury and the magnetic resonance imaging (MRI) was 3.5 years. Exclusion criteria were based on pre-existing developmental disorders, central neurological disorders, intellectual disabilities and musculoskeletal disease. The demographic and clinical descriptors of the TBI group are given in Table 1 and 2.

The study was approved by the Ethics Committee for biomedical research at the KU Leuven and the patients were all recruited from several rehabilitation centres in Belgium (Principal Investigator, Stephan Swinnen). Written informed consent was obtained from either the participants themselves or from the patients’ first-degree relatives, according to the Declaration of Helsinki.

**Table 1:**
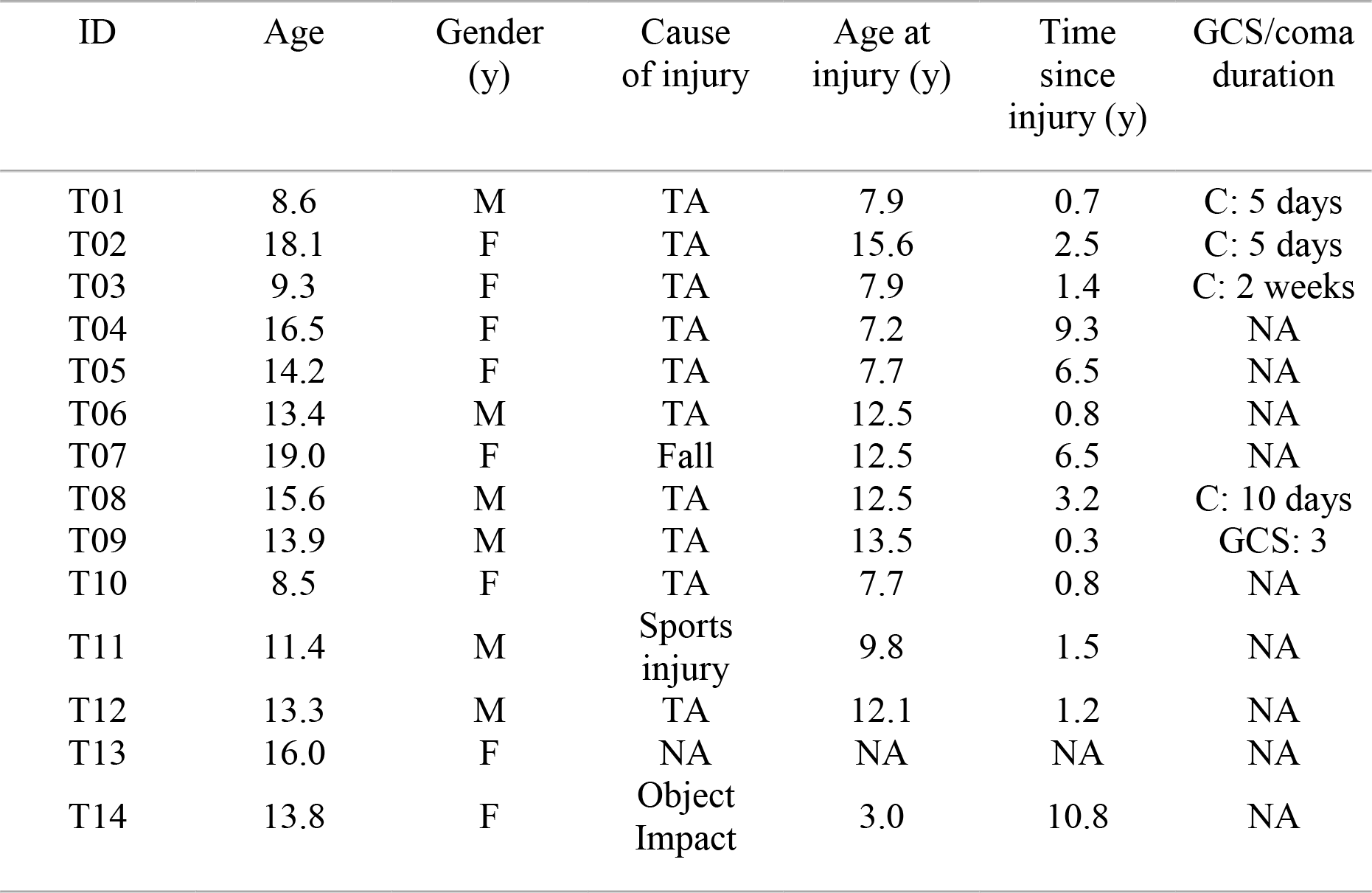
Demographic data of TBI patients. TA = Traffic Accident; C = Coma; NA = Information not available; GCS = Glasgow Coma Scale score; M = male; F = female

| ID | Age | Gender<br>(y) | Cause<br>of injury | Age at<br>injury (y) | Time<br>since<br>injury (y) | GCS/coma<br>duration |
| --- | --- | --- | --- | --- | --- | --- |
| T01 | 8.6 | M | TA | 7.9 | 0.7 | C: 5 days |
| T02 | 18.1 | F | TA | 15.6 | 2.5 | C: 5 days |
| T03 | 9.3 | F | TA | 7.9 | 1.4 | C: 2 weeks |
| T04 | 16.5 | F | TA | 7.2 | 9.3 | NA |
| T05 | 14.2 | F | TA | 7.7 | 6.5 | NA |
| T06 | 13.4 | M | TA | 12.5 | 0.8 | NA |
| T07 | 19.0 | F | Fall | 12.5 | 6.5 | NA |
| T08 | 15.6 | M | TA | 12.5 | 3.2 | C: 10 days |
| T09 | 13.9 | M | TA | 13.5 | 0.3 | GCS: 3 |
| T10 | 8.5 | F | TA | 7.7 | 0.8 | NA |
| T11 | 11.4 | M | Sports<br>injury | 9.8 | 1.5 | NA |
| T12 | 13.3 | M | TA | 12.1 | 1.2 | NA |
| T13 | 16.0 | F | NA | NA | NA | NA |
| T14 | 13.8 | F | Object<br>Impact | 3.0 | 10.8 | NA |

**Table 2:** Clinical data of TBI patients.

| ID | Acute MRI scan within 24 h after injury lesion location/pathology | MRI scan at test session lesion location/pathology |
| --- | --- | --- |
| T01 | Subdural hematoma R FL/PL/TL; cortical contusion R FL/PL; DAI in R FL | Micro-bleeding R centrum semiovale and CC |
| T02 | Subdural hematoma/hemorrhagic contusion TL/FL; injuries R FL, thalamus, R cerebral peduncle, L mesencephalon; cortical and subcortical hemorrhagic areas in PL/TL | Injuries surrounding drain trajectory in RH (superior frontal gyrus, head nucleus caudatus, crus anterior of internal capsule, thalamus, and pons) |
| T03 | DAI in L TL/FL, R TL/FL/PL | Contusion: R anterior temporal pole and R orbitofrontal cortex; Injuries and atrophy in CC (body and splenium); Atrophy of R pons; Hemosiderin deposits in L cerebellar hemisphere, R nucleus entiformis, L/R FL, L/R PL and R PL |
| T04 | Epidural hematoma R FL/TL; shift midline | Injuries in R medial frontal gyrus. Atrophy of the cerebellum; injuries at the level of L FL, premotor cortex, medial frontal gyrus, cingulum, orbitofrontal cortex (L > R); contusion anterior temporal pole (R > L); hemosiderin deposits in CC, L thalamus, striatum (R > L) |
| T05 | NA |  |
| T06 | Hemorrhagic contusion L TL; brain oedema | Hemosiderosis; DAI PL (superior and inferior), R cerebellum, L superior frontal gyrus |
| T07 | Subdural hematoma L FL/TL/PL | Hemosiderin deposits R cerebellar vermis |
| T08 | DAI R TL, internal capsule, supra-orbital R FL, L FL WM (anterior corona radiata), L middle cerebellar peduncle | Atrophy cerebellum; Contusion R FL WM |
| T09 | DAI FL, TL, L OL (hemorrhagic injury), cerebellum, CC, external capsule, R globus pallidus, L thalamus, R cerebral peduncle, R mesencephalon | DAI L FL, periventricular WM, body and genu CC, L thalamus a, R external capsule, anterior TL (L > R), cerebellum; Limited atrophy cerebellum |
| T10 | NA | Enlarged fourth ventricle, atrophy of cerebellar vermis, contusion R cerebellar vermis, hypotrophy of middle cerebellar peduncle and L pons; contusion L TL; hemosiderin deposits R FL, L TL, CC; ventricle drain |
| T11 | Contusion L FL/TL; enlarged, asymmetric ventricle (temporal horn) | DAI splenium CC; ventricular drain |
| T12 | DAI in genu and splenium CC, L FL | Hemosiderin deposits L FL, genu CC |
| T13 | NA | Mild atrophy in cerebellum and cerebrum, more pronounced atrophy in frontal cortices, enlarged ventricles; contusion L/R anterior temporal pole and L/R orbitofrontal cortex. Hemosiderin deposits in cerebellum, R FL |
| T14 | Hemorrhagic contusion L FL, atrophy L FL | Contusion: L anterior middle frontal gyrus and L anterior superior frontal gyrus |

\*WM=white matter; RH=right hemisphere; LH=left hemisphere; FL=frontal lobe; TL=temporal lobe; PL=parietal lobe; OL=occipital lobe; CC=corpus callosum; R = right; L = left. Other codes: y = years.

### 2.2 Behavioural measures of postural control performance

Balance control was assessed using three protocols from the EquiTest System (NeuroCom International, Clackamas, Oregon).

#### 2.2.1 The Sensory Organization Test (SOT)

This test measures static postural control while subjects are standing as still as possible, barefoot, on a movable platform (forceplate) under 4 sensory conditions: 1) eyes open, fixed platform; 2) eyes closed, fixed platform; 3) eyes open with the platform tilting in response to body sway to prevent the ankles from bending (reduced somatosensory feedback); 4) eyes closed, tilting platform. In order to familiarize the subject with the test and to avoid any initial effect of surprise on the sensory manipulations, we included 1 practice trial for each condition before completing the actual measurements. After that, each condition was repeated three times in a randomized order. Each trial lasted twenty seconds. We used an established protocol applied in earlier studies to assess balance control in young and older healthy adults, calculating the centre of pressure (COP) trajectory from the forceplate recordings (100 Hz) (Van Impe *et al.*, 2013). A mean SOT balance score was acquired for each condition from the 3 trials, excluding trials in which the subject fell. We evaluated the behavioural outcome through the inverse path-length (iPL) of the COP trajectory to acquire a SOT balance index in which higher scores reflect better balance control (and less body sway).

#### 2.2.2 The Limits Of Stability Test (LOST)

This is a more dynamic test of balance control that involves goal-directed postural adjustments, where subjects intentionally displace their centre of gravity (COG) in different directions without stepping, falling or lifting their heel or toes. At the beginning of each trial, the COG (provided by the Equitest forceplate) was positioned in the centre, as indicated by a representation on a screen in front of the subject. On presentation of a visual cue and by leaning over in the right direction, the subject had to move the COG from the center towards one of the radial targets presented on the screen as quickly and accurately as possible. The following eight target directions were assessed: front, right front, right, right back, back, left back, left, and left front. After two practice trials, each direction was assessed once in a random order. The trial was interrupted and repeated if the subject fell or took a step, and that trial was not analyzed. Directional control (DC) was computed as the outcome measure reflecting dynamic balance control. Specifically, the DC (expressed as a percentage) was calculated as the difference between on-target (in target direction) and off-target movement (extraneous movement) divided by the amount of on-target movement, as follows: (amount of on-target movement - amount of off-target movement)/(amount of on-target movement) × 100%. Higher scores reflect better DC and only a straight line towards the target would result in a score of 100%, with no off-target movements. Finally, to end up with a single measure to be correlated with imaging results, DC scores were averaged across the eight target directions for further analysis.

#### 2.2.3 The Rhythmic Weight Shift Test (RWST)

Like the LOST, this is a dynamic test of balance control measuring the ability to move the COG rhythmically from right to left, or forward and backwards, between two target positions. Each direction (backward-forward, left-right) was performed at 3 different speeds: slow (a pace of 3 seconds between each target), medium (a pace of 2 seconds) and fast (a pace of 1 second). Each combination of speed and direction (a total of 6 combinations) was performed in a separate trial of 6 movement repetitions that were preceded by 4 practice repetitions. The trial was interrupted and repeated if the subject fell or took a step. The DC was calculated as above (similar to LOST) and DC scores were averaged across directions and velocities for further analysis.

In summary, postural control was evaluated through three different score indexes: one measuring static postural control (iPL-SOT), and two measuring dynamical postural control (DC-LOST and DC-RWST). The three indexes were used as behavioural outcome to correlate with imaging results.

### 2.3. MRI data acquisition

MRI scanning was performed on a Siemens 3T Magnetom Trio MRI scanner with a 12- channel matrix head coil.

#### 2.3.1. Anatomical data

A high resolution T1 image was acquired with a 3D magnetization prepared rapid acquisition gradient echo (MPRAGE): repetition time [TR] = 2300 ms, echo time [TE] = 2.98 ms, voxel size = 1 × 1 × 1.1mm^3^, slice thickness = 1.1 mm, field of view [FOV] = 256 × 240mm^2^, 160 contiguous sagittal slices covering the whole brain and brainstem).

#### 2.3.2. Diffusion Tensor Imaging (DTI)

A DTI SE-EPI (diffusion weighted single shot spin-echo echoplanar imaging) sequence was acquired with the following parameters: [TR] = 8000 ms, [TE] = 91 ms, voxel size = 2.2 × 2.2 × 2.2 mm^3^, slice thickness = 2.2 mm, [FOV] = 212 × 212 mm^2^, 60 contiguous sagittal slices covering the entire brain and brainstem. A diffusion gradient was applied along 64 non-collinear directions with a b-value of 1000 s/mm^2^. Additionally, one set of images was acquired with no diffusion weighting (b=0 s/mm^2^).

#### 2.3.3. Resting state functional data

Resting state fMRI time series were acquired over a 10 minute session using the following parameters: 200 whole-brain gradient echo echoplanar images with TR/TE = 3000/30 ms; FOV = 230 × 230mm^2^; voxel size = 2.5 × 2.5 × 3.1mm^3^, 80 × 80 matrix; slice thickness = 2.8 mm; 50 sagittal slices, interleaved in descending order.

### 2.4. MRI data pre-processing

#### 2.4.1. Diffusion Tensor Imaging

The DTI data were pre-processed using FSL (FMRIB Software Library v5.0) and the Diffusion Toolkit (Wang *et al.*, 2013). An eddy current correction was applied to overcome the artefacts produced by variation in the direction of the gradient fields of the MR scanner, together with the artefacts produced by head movements. Using the corrected data, a local fitting of the diffusion tensor was applied to compute the diffusion tensor model for each voxel. Next, a Fiber Assignment by Continuous Tracking (FACT) algorithm was applied (Mori *et al.*, 1999). We then computed the transformation from the Montreal Neurological Institute (MNI) space to the individual-subject diffusion space and projected a high resolution functional partition to the latter, composed by 2,514 regions of interest (ROIs) and generated after applying spatially constrained clustering to the functional data using the method explained in (Craddock *et al.*, 2012). This allowed a 2,514 × 2,514 structural connectivity (SC) matrix to be built for each subject by counting the number of white matter streamlines connecting all ROI pairs within the 2,514 regions partitioned. Thus, the element matrix (i,j) of SC is given by the streamlines number between regions i and j. SC is a symmetric matrix, where connectivity from i to j is equal to that from j to i. Finally, we made the SC matrices binary for the analysis, considering only two possible values: 0 when no streamlines existed between i and j; and 1, when any non-zero number existed between the two regions i and j.

#### 2.4.2. Resting state functional MRI

The fMRI data was pre-processed with FSL and AFNI (http://afni.nimh.nih.gov/afni/). In the first place, the fMRI dataset was aligned to the middle volume to correct for head movement artefacts and slice-time correction was then applied for temporal slice-alignment. All voxels were spatially smoothed with a 6 mm full width at half maximum (FWHM) isotropic Gaussian kernel and after intensity normalization, a band pass filter was applied between 0.01 and 0.08 Hz (Cordes *et al.*, 2012), followed by the removal of linear and quadratic trends. We next regressed out the movement time courses, the average cerebrospinal fluid (CSF) signal, the average white-matter signal and the average global signal. Finally, the functional data was spatially normalized to the MNI152 brain template, with a voxel size of 3^*^3^*^3 mm^3^.

### 2.5. Clustering regions of interest into modules by using a new hierarchical brain atlas

The initial 2,514 ROIs were grouped into modules using a recently published atlas (Diez *et al.*, 2015) in which the regions are functionally coherent (i.e. the dynamics of voxels belonging to one region are very similar) and at the same time, they are structurally wired (i.e. the voxels belonging to a given region are interconnected through white matter fibers). Some existing atlases are purely anatomical or structural (Lancaster *et al.*, 2000; Tzourio-Mazoyer *et al.*, 2002; Eickhoff *et al.*, 2005; Desikan *et al.*, 2006), and others are purely functional, like the well-known Brodmann atlas based on lesion studies or data-driven methods (Craddock *et al.*, 2012). Although obtaining suitable brain partitions (or atlases) has been studied intensively (see Craddock *et al.*, 2013 and references therein), to the best of our knowledge, we were the first to propose a brain partition that accounts for regions that are relevant to both structure and function (Diez *et al.*, 2015) which is now implemented in the current project.

Although full details are given in (Diez *et al.*, 2015), here, we briefly summarize the hierarchical clustering approach which, applied to a combination of functional and structural datasets, results in a hierarchical tree or dendrogram in which nodes were progressively merged together into M different moduli following a nested hierarchy of “similarity” (which reflects correlation for the functional data and white matter streamlines-number for the structural one). Thus, cutting the tree at a certain level led to a pooling of the initial 2,514 ROIs into a finite number of modules 1 ≤ M ≤ 2,514 (in principle, an arbitrary number for M can be obtained by varying the depth of the cut). Thus, to provide some examples, the highest dendrogram level M=1 corresponded to all 2,514 regions belonging to a single module, coincident with the entire brain, whereas the lowest level M=2,514 corresponded to 2,514 separated modules, all of them composed of one single ROI.

It was also shown (Diez *et al.*, 2015) that the hierarchical brain partition with M=20 modules was optimal based on cross-modularity, an index simultaneously accounting for three features: 1) Modularity of the structural partition; 2) Modularity of the functional partition; and 3) Similarity between functional and structural modules. The matlab code to calculate the cross-modality index between structural and functional connectivity matrices can be downloaded at http://www.nitrc.org/projects/biocrhcatlas/

To compute cross-modularity, we first assessed modularity simply for accounting the quality of the brain partition, i.e., a partition with high modularity has modules highly isolated from each other, achieved for instance by maximizing the fraction of intra-module to inter-module connections with respect to randomizations. In particular, we applied the Newman’s algorithm (Newman, 2004) to address modularity. In addition to modularity, cross-modularity made use of the similarity between structural and functional modules, which was approached by calculating the Sorensen’s index (Sorensen, 1948), a normalized quantity equal to twice the number of common connections in the two partitions divided by the total number of connections in the two partitions.

The entire hierarchical brain partition can be downloaded at http://www.nitrc.org/projects/biocrhcatlas/

### 2.6 Group differences in structural networks

Between the regions defined from the hierarchical atlas, structural networks (SN) were assessed by counting all the connections (ie. streamlines) starting from one of the regions and ending in a different one. Notice that regions can be defined at any level of the hierarchical tree, cf. section 2.5. We then calculated the region’s connectivity degree: the total number of connections reaching a region (which coincides with the total number of connections leaving it, as the structural connectivity is a symmetric matrix). A one-way ANOVA test was then applied to these connectivity maps to search for significant differences (*p* < 0.05).

To assess the significance of the structural differences, we applied a permutation test by performing 1,000 random permutations of the 2,514 matrix, calculating the region’s degree for each permutation and keeping the same size for all 2,514 regions. We then generated the probability distribution for these values, which constitutes the null-hypothesis since all the dependencies have been removed by the shuffling procedure. All regions with p > 0.05 were discarded.

### 2.7 Group differences in resting state brain dynamics within individual regions

In order to measure group differences in the resting state brain dynamics within each of the M=20 regions, we first obtained the time-series of the first principal component for each region, chosen as the best representative for the entire region. Next, we compared 4 different descriptors extracted from these time series: variance (2^nd^ standardized moment; quantifies fluctuation size), skewness (3^rd^ standardized moment; identifies extreme brain dynamics in the resting state (Amor *et al.*, 2015), as it measures how much asymmetry a distribution has with respect to its mean), kurtosis (4^th^ standardized moment; measures the long-tail effect on the data distribution), and the number of points resulting from the point process analysis (PPA) (Tagliazucchi *et al.*, 2012) (measured by counting the number of amplitude peaks in the BOLD signal, and in particular, counting the points with value greater than the mean value of the time series plus 1 SD). Previously, PPA has been successfully used (Tagliazucchi *et al.*, 2012) to encode relevant information of the resting-state fMRI into discrete events, precisely corresponding to the large-amplitude signal-peaks quantified by PPA. These descriptors were subjected to a one-way ANOVA test to evaluate the differences between the TBI and control participants (p <0.05).

### 2.8 Group differences in functional networks

Motivated by an earlier study (Smith *et al.*, 2009), functional networks (FN) were assessed by quantifying the interaction between each of the M=20 regions and the rest of the brain (Supplementary Fig. S1). First, within each of the M=20 regions, we applied a principal component analysis (PCA) to reduce the dimension of the data, resulting in 20 components for each of the M=20 regions. Next, we applied an independent component analysis (ICA) to obtain C=20 independent components associated to each of the M=20 regions. Finally, we applied a spatial regression method to quantify the contribution of each brain voxel to each component (i.e., component’s spatial map). Next, we clustered all spatial maps by applying the k-means clustering-algorithm using the spatial correlation between observations as the similarity measure (Bishop, 2006); thus, two maps were considered to belong to the same cluster if they showed high spatial correlation. After k-means, the 820 observations per region (41 subjects, C=20 independent components) were grouped into 5 clusters, that we named the 5 most representative clusters (MRCs). Here, the number 5 was chosen by careful inspection to guarantee a good discrimination between the different clusters. K-means, in addition to return the 5 MRCs, also provides one label for each of the 820 observations, 1,2,3,4 or 5, indicating to which MRC the observation belongs.

In order to compare groups, control vs. TBI, we applied a t-test between observations belonging to the same MRC.

For multiple comparison correction, due to a voxel-by-voxel analysis, a statistical significant cluster-wise correction was applied. Monte-Carlo simulation (3dClustSim, AFNI, http://afni.nimh.nih.gov) was performed with 10,000 iterations to estimate the probability of false positive clusters with a p value <0.05 in the analysis. After correcting for multiple comparisons, three classes of activity maps for each region were calculated: 1) the average FN in control participants (corresponding to the contrast [1, 0]); 2) the average FN in TBI (contrast [0, 1]); and 3) the differences in average FN between control and TBI (applying the two different contrasts [1, −1] and [−1, 1] we achieve, respectively, control > TBI connectivity and TBI > control connectivity).

## 3. Results

### 3.1 TBI-induced alterations in postural control performance

Alterations in postural control performance were measured by three different tests (Methods). First, the Sensory Organization Test (SOT) showed that TBI patients had a smaller inverse path-length (iPL) in the COP trajectory as compared to control participants (Control: 106.72 +- 8.41; TBI: 88.63 +- 28.80; *p* = 0.0041), which reflects that TBI patients had a worse balance (more body sway) than control. Analogously, the Rhythmic Weight Shift test (RWST) showed that TBI patients performed worst as compared to control (control: 84.20 +- 6.66; TBI: 77.11 +- 9.82; p = 0.0073), again, confirming a poorer dynamic balance for TBI patients. In contrast, the limits of stability test (LOST) did not show any difference between groups with respect to the dynamic control (DC) index (Control: 84.31 +- 6.44; TBI: 83.46 +- 5.89; *p* = 0.6776).

### 3.2. TBI-induced alterations in structural networks

Alterations in structural networks were assessed by calculating the connectivity degree for each region in the hierarchical atlas from the inter-region connectivity matrix and performing a group comparison (after correcting for multiple comparisons by performing random permutations in the structural connectivity matrix). For all M=20 regions (Table 3) control participants showed a higher number of connections reaching a region than TBI patients (Fig. 1). This suggests a global decrease in connectivity associated with TBI. More specifically, significant differences in connectivity degree were evident within region 6 (including part of the cerebellum, lateral occipital sulcus, fusiform gyrus, occipital-temporal junction and superior parietal gyrus), region 14 (including part of the hippocampus and parahippocampal gyrus, amygdala, putamen, insula, ventral diencephalon, temporal gyrus and temporal pole), region 18 (including part of the hippocampus and entorhinal cortex, fusiform gyrus, inferior and middle temporal gyrus and parahippocampal gyrus), and region 20 (including part of the cerebellum and parahippocampal gyrus).

**Fig. 1:**
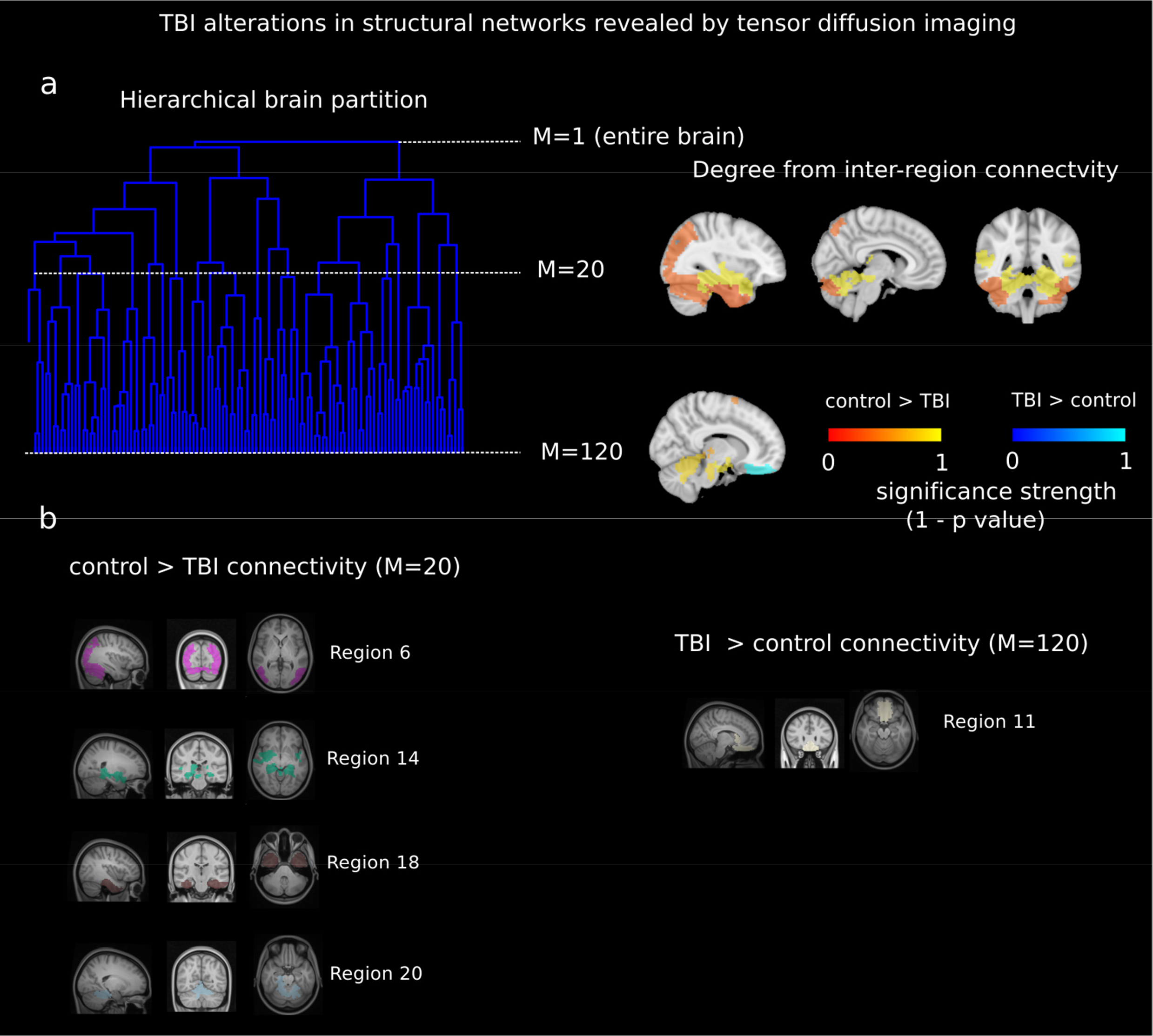
TBI-induced alterations to structural networks revealed by diffusion tensor imaging. **a:** Hierarchical tree or dendrogram defining the brain partition published in [22] where three different levels of the tree have been emphasized: M=1, where all brain regions belong to a single module; M=20, the optimal brain partition (see Methods); and M=120, the level at which structural connectivity was higher in TBI than in the controls. Group differences were calculated on region degree maps calculated on the inter-region connectivity matrix and after a one-way ANOVA test (p < 0.05). Multiple comparison corrections were achieved by applying random permutations to the 2,514 matrix and calculating the region’s degree of connectivity for each, thereby building the null-hypothesis distribution as all correlations were removed by shuffling. Greater connectivity in controls than in TBI (red scale) was found at M=20 for all brain regions and at M=120, where a region with TBI > control connectivity was also found (blue scale). **b:** At M=20 (left graph), significant control; > TBI connectivity was evident in: regions 6 (including part of the cerebellum, lateral occipital sulcus, fusiform gyrus, occipital-temporal junction and superior parietal gyrus); region 14 (including part of the hippocampus and parahippocampal gyrus, amygdala, putamen, insula, ventral diencephalon, temporal gyrus and temporal pole); region 18 (including part of the hippocampus and entorhinal cortex, fusiform gyrus, inferior and middle temporal gyrus and parahippocampal gyrus); and region 20 (including part of the cerebellum and parahippocampal gyrus). At M=120 (right graph), TBI > control connectivity was found within the region 11, including part of the rectus and the superior and inferior frontal orbital gyrus.

**Table 3:**
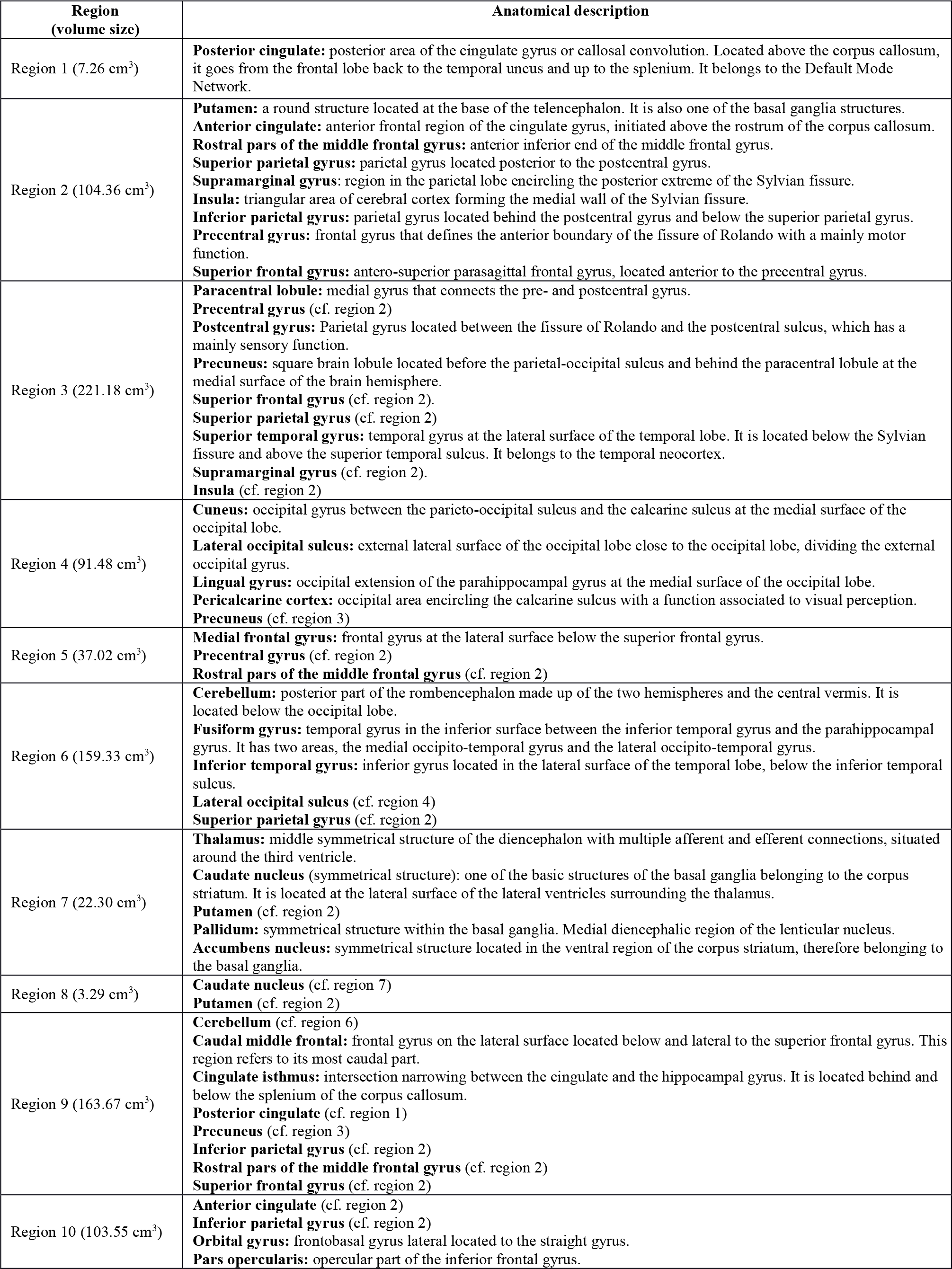
Anatomical description of the M=20 regions in the hierarchical atlas published in [22]. In the first column, we also indicate the volume of the region.

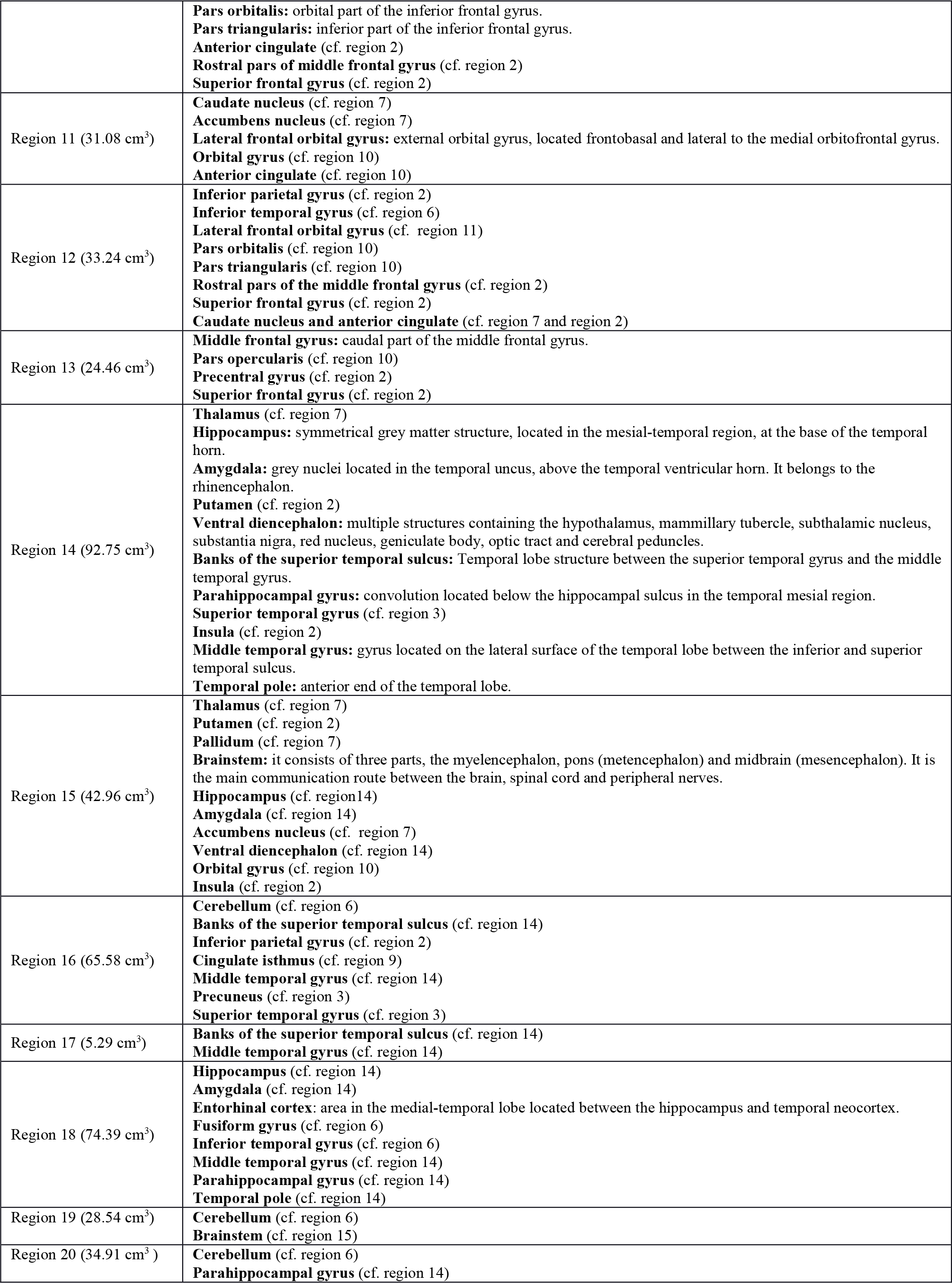

At the level of M=20 in the hierarchical tree the inter-region connectivity degree was higher in control for all the regions, indicating that one needs to go down through the hierarchical tree to find another representation with a higher spatial scale (the number of M regions is smaller as the tree goes down) and where the TBI connectivity can be higher as compared to control. Proceeding in this way, we found at the level of M=120 regions that TBI had bigger connectivity than control within the region 11 of the hierarchical atlas, a region including part of the rectus and the superior-inferior orbital frontal.

### 3.3 TBI-induced alterations in resting state brain dynamics within individual regions

Alterations in the resting state brain dynamics within each of the M=20 regions were assessed by calculating the differences in the time series of the first principal component extracted from each region (Fig. 2). In particular, differences were addressed with respect to the second moment (i.e. the variance), the third moment (skewness), the forth moment (kurtosis) and the number of time-series points that had a value above the mean value of the time series plus 1 times the standard deviation (the so-called Point Process Analysis, PPA [41]). After repeating the same procedure for all M=20 regions of the hierarchical atlas, significant differences were only present within regions 10 and 11, and for the two regions, the TBI brain dynamics descriptors were higher than in the control group. In particular, TBI > control variance was found in region 11, TBI > control kurtosis was found in region 10, and the number of points after the PPA bigger in TBI was found in region 11, but no differences were found when comparing the skewness (Fig. 2). Region 10 roughly consisted of the frontal part of the default mode network, including part of the anterior cingulate gyrus, orbital gyrus, pars opercularis, pars orbitalis, pars triangularis, rostral pars of the middle frontal gyrus and superior frontal gyrus. Region 11 consisted in part of the caudate nucleus, nucleus accumbens, lateral frontal orbital gyrus, orbital gyrus and anterior cingulate gyrus.

**Fig. 2:**
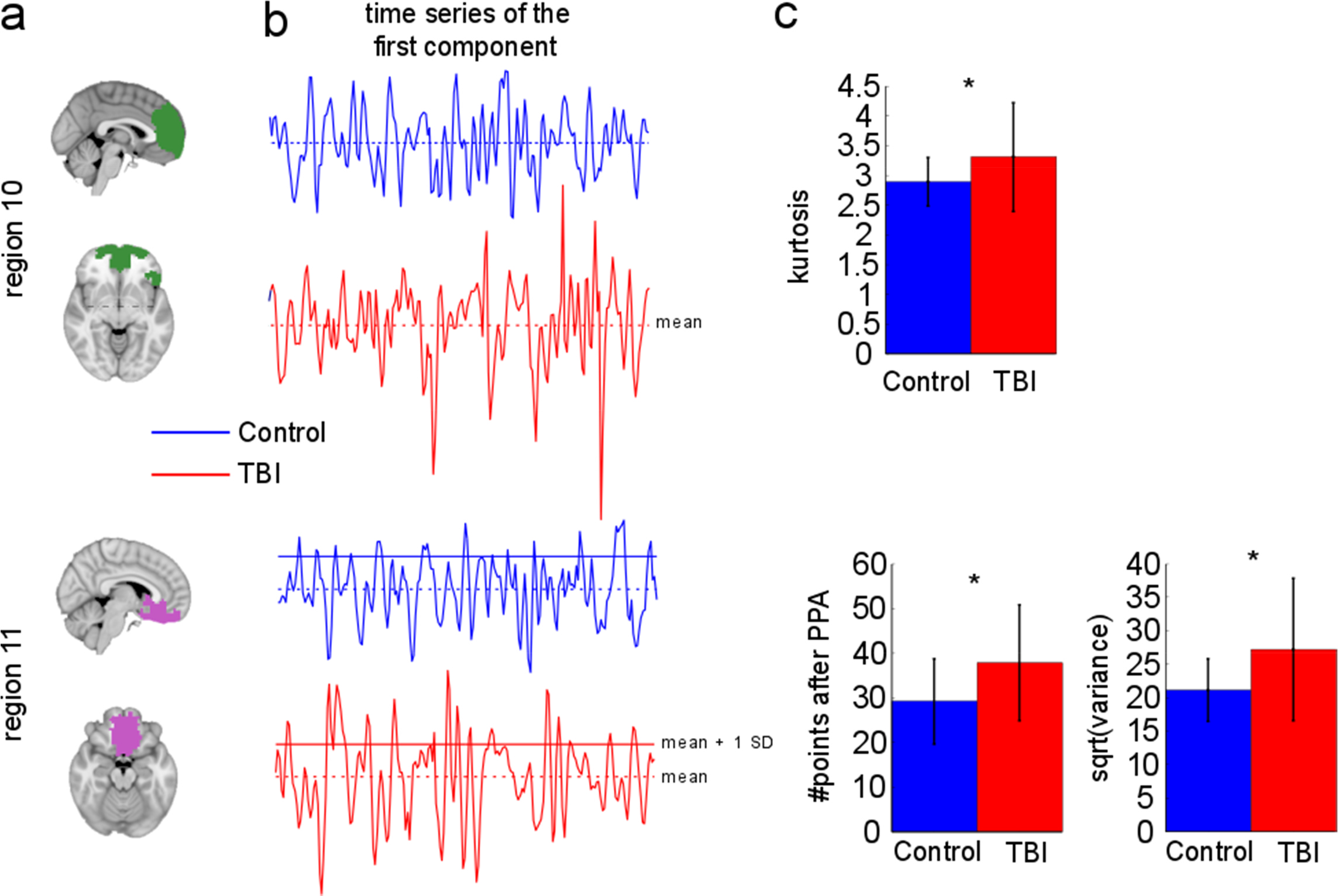
TBI-induced alterations to brain dynamics within individual regions revealed by resting state fMRI. For each of the M=20 regions in the hierarchical atlas, we extracted the time-series of the first principal component and calculated four different descriptors: 1) the variance; 2) skewness; 3) kurtosis; and 4) the number of points after the PPA (see Methods). **a:** Only two regions showed differences between the TBI and control, regions 10 and 11, both localized in the frontal lobe of the brain. **b:** Time series of the first principal component. The dashed lines represent the mean value of the time series and the solid lines represent the threshold used for the PPA, here equal to the mean + 1 SD. c: For region 10, only kurtosis showed differences between the TBI and control subjects. For region 11, the number of points after the PPA and the variance of the first component (plotted here as its square root, i.e.: the standard deviation) differed between TBI and control subjects. For both regions 10 and 11 the time series of the first component showed compensation rather than a deficit, i.e.: TBI activity (red bars) was higher than the controls (blue bars).

### 3.4 TBI-induced alterations in functional networks

Functional networks were addressed by quantifying the interaction from each of the M=20 regions to the rest of the brain. Within each region, we first obtained C=20 components (after PCA followed by ICA) and next, we performed spatial regression of the C=20 PCA+ICA components to all the brain voxels, in this way obtaining C=20 spatial maps for each of the regions. Three important issues arose in relation to this procedure: 1) several components belonging to different regions provided the same spatial map; 2) the spatial maps derived from each component were noisy; and 3) the inter-subject variability was high. As explained in methods, we overcame these limitations by grouping all the spatial maps with a similar spatial distribution into clusters with the k-means algorithm, grouping the 820 observations (41 subjects C=20 independent components) per each region, into the 5 most representative clusters (MRCs).

After this procedure, it was possible to obtain the same MRC from different regions. In particular, Fig. 3 show the results associated to one of the MRCs obtained from the following regions: region 3 (including part of the sensory-motor and auditory networks), regions 14 and 15 (including part of the thalamus, hippocampus, amygdala, putamen, ventral diencephalon and insula), region 18 (including part of the hippocampus and entorhinal cortex, fusiform gyrus, inferior and middle temporal gyrus and parahippocampal gyrus), region 19 (including part of the cerebellum and brainstem) and region 20 (including part of the cerebellum and parahippocampal gyrus).

**Fig. 3:**
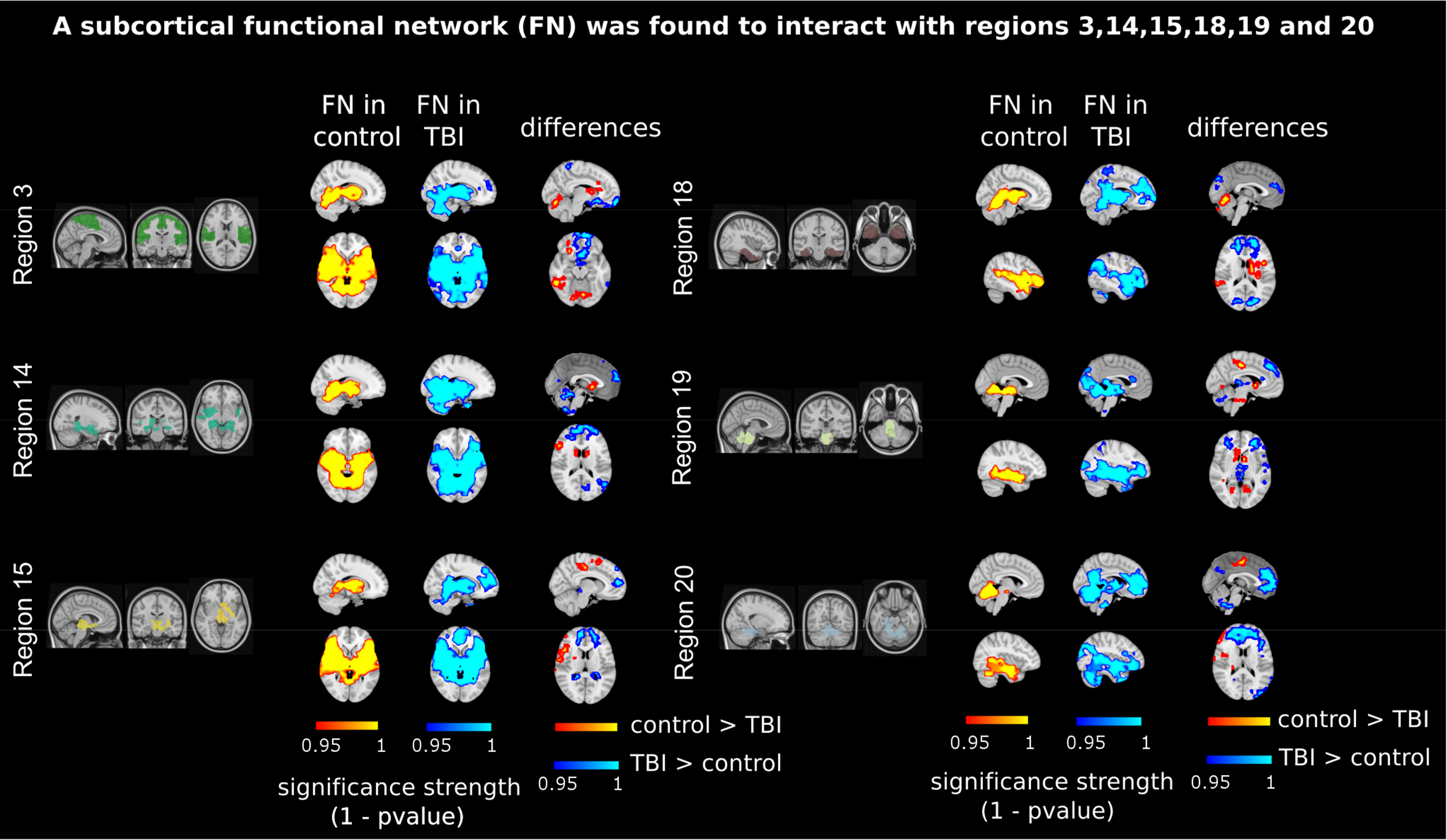
Prefrontal activation at rest triggered by subcortical interactions. Significant regions after using different contrasts: Column 1,”FN in control”, with a red bar and corresponding to the contrast [1, 0] (Methods); Column 2,”FN in TBI”, with a blue bar and corresponding to the contrast [0, 1]; Column 3,”differences” and corresponding to two contrasts [1, −1] and [−1, 1] represented in red (control > TBI activation) and blue (TBI > control activation), respectively. TBI patients (but not control participants) incorporated to the subcortical network also the prefrontal part of the brain (coloured in blue at the column "differences"). In all cases, the bar-scale represents the significance strength, measured by 1 minus the p value. Contrasts [1, 0] and [0, 1] define the “subcortical network" (which corresponds to one MRC including part of the cerebellum, the basal ganglia, the thalamus, the amygdala and the temporal poles). This network resulted from the interactions coming from region 3 (including part of the sensory-motor and auditory networks), regions 14 and 15 (including part of the thalamus, hippocampus, amygdala, putamen, ventral diencephalon and insula); region 18 (including part of the hippocampus and entorhinal cortex, fusiform gyrus, inferior and middle temporal gyrus, and parahippocampal gyrus); region 19 (including part of the cerebellum and brainstem); and region 20 (including part of the cerebellum and parahippocampal gyrus).

The anatomical representation of this MRC (obtained from regions 3, 14, 15, 18, 19 and 20) revealed that it was composed of a *subcortical network* (see Fig. 3 - columns “FN in control” and “FN in TBI”), consisting of part of the motor network, basal ganglia, cerebellum, thalamus, parahippocampus, hippocampus, precuneus, amygdala, insula, caudate nucleus, putamen and pallidum. Interestingly, TBI > control connectivity (obtained with the contrast [−1, 1]) was found in the frontal lobe (Fig. 3 - column “differences”, colored in blue), within a region consisting of part of the middle frontal and superior orbital gyrus, rectus, olfactory lobe, frontal medial orbital, precuneus and cingulum anterior. In other words, the subcortical network illustrated in Fig. 3 with labels “FN in control” and “FN in TBI” included the prefrontal brain in TBI patients but this did not happen for control participants.

Another MRC resembled the *task-positive network* (Fig. 4 - columns “FN in control” and “FN in TBI”). In particular, this MRC consisted of parts of the cerebellum, lingual gyrus, fusiform gyrus, inferior occipital gyrus, calcarine sulcus, cuneus, precuneus, superior temporal pole, superior motor area and insula. This MRC was the result of functional interactions affecting region 1 (including part of the posterior cingulate gyrus), region 4 (the medial visual network), region 5 (including part of the medial frontal gyrus, precentral gyrus and rostral pars of the middle frontal gyrus), region 12 (including part of the inferior parietal gyrus, inferior temporal gyrus, lateral frontal orbital gyrus, pars orbitalis, pars triangularis, rostral pars of the middle frontal gyrus, superior frontal gyrus, caudate nucleus and anterior cingulate gyrus), and regions 14 and 15 (including part of the thalamus, hippocampus, amygdala, putamen, ventral diencephalon and insula).

**Fig. 4:**
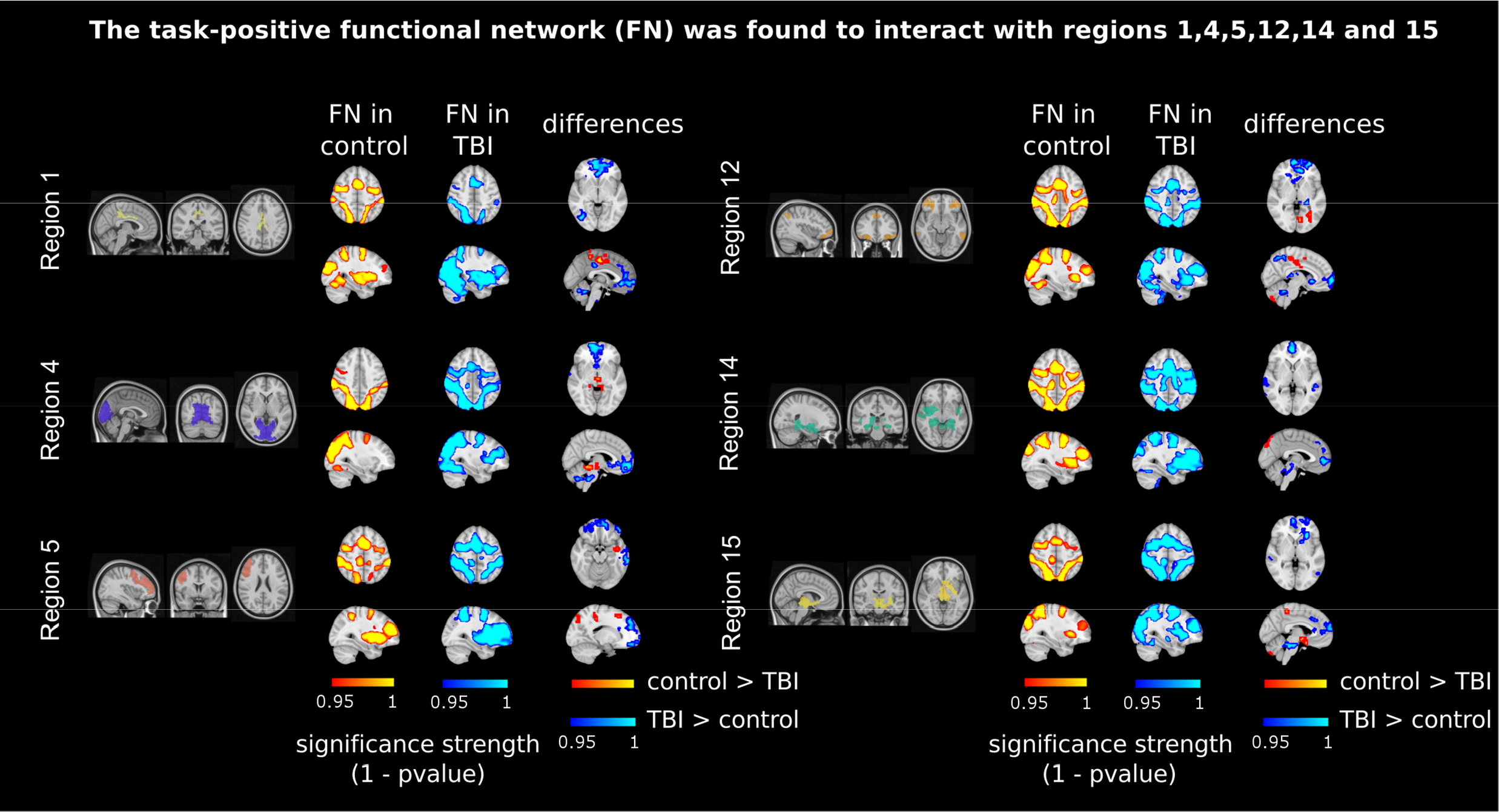
Prefrontal activation at rest triggered by task-positive interactions. As in Fig. 3 but the MRC now resembles the task-positive network (see labels "FN in control" and "FN in TBI"), which is now resulting from region 1 (posterior cingulate cortex); region 4 (medial visual cortex); region 5 (medial frontal gyrus); region 12 (inferior parietal and temporal gyrus, lateral frontal orbital gyrus, rostral pars of the middle frontal gyrus and pars orbitalis and triangularis); and regions 14 and 15 (subcortical structures). Similar to what it was shown in Fig. 3, now TBI patients incorporated to the task-positive network the prefrontal part of the brain (coloured in blue at the column "differences").

Although both control and TBI groups provided a similar task-positive network (Fig. 4 - columns “FN in control” and “FN in TBI”), however, TBI > control connectivity (Fig. 4 - column “differences”, colored in blue) was found in the frontal brain, and in particular, within a region consisting of part of the frontal medial orbital, anterior cingulum, precuneus, superior frontal and angular gyrus. Thus, similar to what happened for the subcortical network represented in Fig. 3, the task-positive network included the prefrontal cortex in TBI patients but did not for control participants.

### 3.5 Brain regions showing increased connectivity in TBI for both functional and structural networks

Since we observed an increased connectivity in TBI patients relative to controls for both functional and structural networks, we decided to take a closer look at these overlapping findings by superimposing these regions (Fig. 5). With regard to the analysis performed for <u>structural networks</u>, a higher connectivity degree in TBI was found in a small subnetwork consisting of a hub that is connected to other regions. The region’s hub (Fig. 5a - plotted in red) belongs to region 11 of the hierarchical atlas and connects to superior frontal regions, anterior cingulum, thalamus, striatum, insula, amygdala, hippocampus and parahippocampus, olfactory lobe and cerebellum (Fig. 5A - areas in green). With regard to the analysis performed for <u>functional networks</u> (Fig. 5B), two regions showed increased connectivity in TBI as compared to controls: one region interacting with a subcortical network (including superior frontal gyrus, superior medial frontal gyrus and middle frontal gyrus, and anterior cingulum) and another region interacting with the task-positive network (including anterior cingulum, frontal medial gyrus, middle orbital gyrus superior frontal medial gyrus and rectus). The overlapping percentage (as measured as the Sorensen’s index between the two networks) between the structural network represented in Fig. 5A and the functional ones represented in Fig. 5B was of 45% for the subcortical network and of 20% for the task-positive network.

**Fig. 5:**
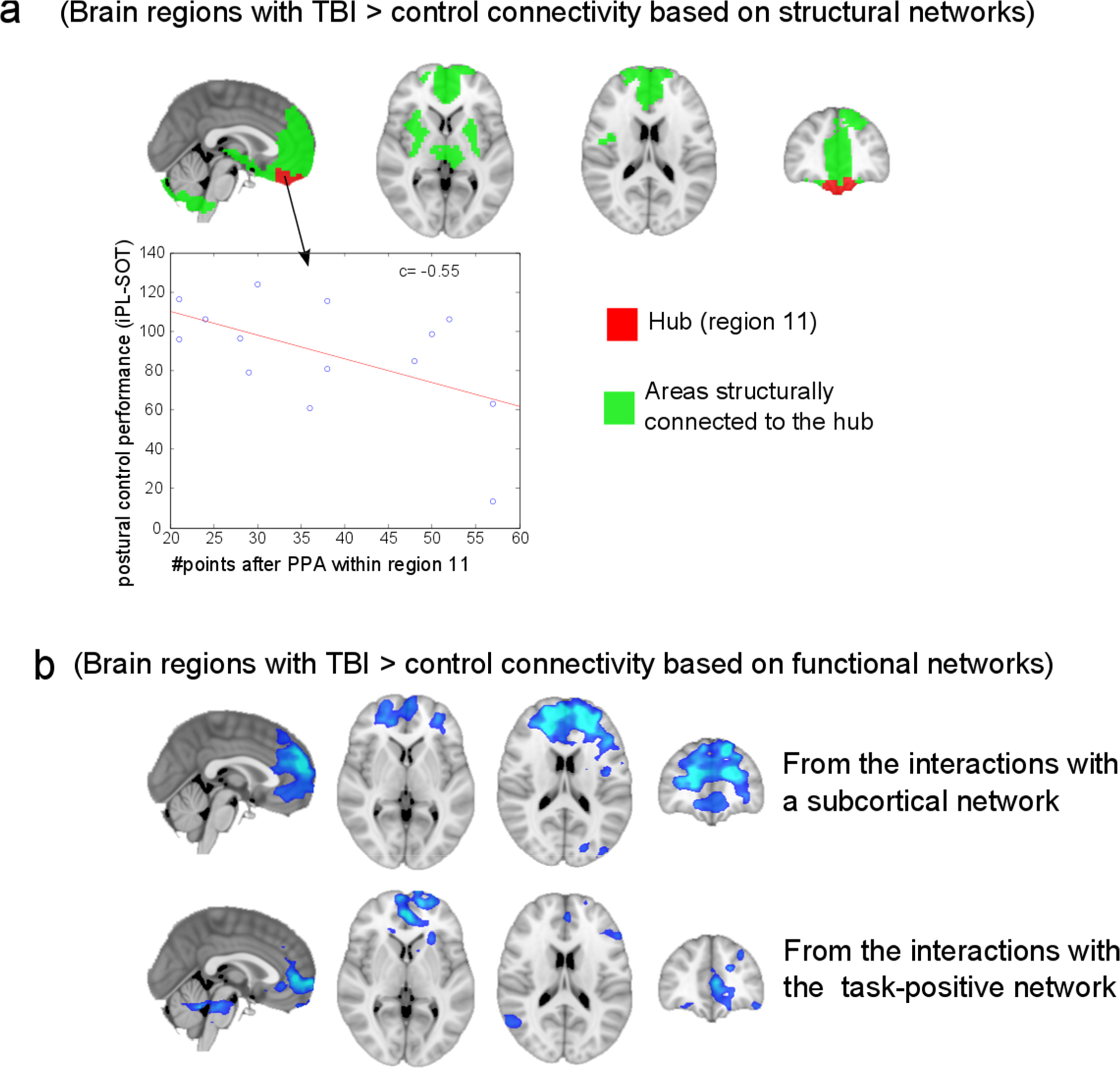
Common regions where TBI > control connectivity occurred for both structural and functional network analyses. **a:** Structural network compensation. TBI > control structural connectivity occurred within a subnetwork consisting of a hub (coloured in red) connected to other regions (coloured in green); the hub includes the orbitofrontal and rectus regions and belongs to the region 11 in the M=20 hierarchical atlas. The regions connected to the hub are frontal superior regions, the anterior cingulum, the thalamus, the striatum, the insula, the amygdala, the hippocampus and para-hippocampus, the olfactory cortex and the cerebellum. The number of points after the Point-Process Analysis (PPA) within region 11 was also correlated with postural control measures; here it is represented the iPL-SOT score. **b:** Functional network compensation. TBI > control functional connectivity (blue) occurred when interacting with subcortical structures (including the superior frontal gyrus, the superior medial frontal and the middle frontal gyri, and the anterior cingulum) and the task-positive network (including the anterior cingulum, the medial frontal and middle orbital gyri, the superior frontal medial gyrus and the rectus). For both situations, the spatial maps represent the functional the result of averaging all spatial maps with contrast [−1, 1] in Fig. 3 (“subcortical network”) and Fig. 4 (“task-positive network”). Thus, TBI patients (but not control participants) incorporated to both the subcortical and the task-positive networks the prefrontal brain of the brain.

### 3.6 Relation between postural control and prefrontal activation at rest

We found that within region 11 of the hierarchical atlas, there was an increase in connectivity for both structural and functional networks for TBI patients when compared to control participants. Correlational analyses revealed that the prefrontal activation at rest of region 11, represented by a high number of points after PPA, was related to the inverse path-length score of the static SOT test (iPL-SOT; *r* = −0.55, *p* = 0.04) and the directional control score of the dynamic test (DC-RWS; *r* = −0.57, *p* = 0.03). Moreover, a tendency towards a significant relationship with DC-LOS was found (*r* = −0.52, *p* = 0.06). These results suggest that a better balance performance is associated with decreased activation in region 11.

Motivated by recent results (Tagliazucchi1 *et al.*, 2014), we chose to correlate (as a good descriptor for the brain dynamics within a region) the number of points after the PPA within region 11 to the three different indices: iPL-SOT, DC-LOS and DC-RWS (Fig. 5A illustrates the results for iPL-SOT, but similar plots can be obtained for DC-LOS and DC-RWS).

## 4. Discussion

Here, we provide the first evidence that TBI-induced alterations in functional and structural networks show overlapping results. With respect to both neuronal networks TBI patients demonstrate increased connectivity as compared with controls within a prefrontal region. Moreover, these TBI-induced network alterations are associated with changes in balance performance.

**TBI-induced alterations in postural control performance.** Alterations in postural control performance were measured by three different tests: SOT, RWST and LOST. We found that TBI patients had a worse balance (more body sway) than control participants.

**TBI-induced alterations in structural networks.** In agreement with previous studies (for reviews see Gentry *et al.*, 1988; Zappala *et al.*, 2012; Hulkower *et al.*, 2013), we found that TBI patients showed reduced structural connectivity (i.e., smaller connectivity degree) for most brain areas compared to healthy participants. We found a strong decrease in connectivity degree in motor areas, brainstem, cingulum, cerebellum and temporal poles, areas that are typically associated with the performance of motor skills and balance control. Indeed, decreased subcortical connectivity, in particular in the brainstem and cerebellum, was recently associated with postural impairments in TBI patients (Drijkoningen *et al.*, 2015b), suggesting a possible diffuse pathology across subcortical structures.

Although for most brain areas we found a lower connectivity degree in TBI relative to controls, we also found a higher connectivity degree in TBI relative to controls exclusively in the prefrontal cortex. This finding, together with the observation that TBI patients have a poorer performance in postural control, may provide a mechanism for stronger cognitive control of such actions.

**TBI-induced alterations in functional networks.** Our approach, focusing on interactions between the regions defined by the hierarchical atlas and the rest of the brain, revealed that TBI patients incorporated the prefrontal cortex together with a subcortical functional network. This result possibly suggests a mechanism compensating for TBI-induced subcortical-cortical axonal disruptions, as confirmed by the results found in the analysis of structural networks, showing decreased white matter connectivity in cortical to subcortical pathways. This disconnection is also consistent with grey matter deficits reported in the frontal and temporal cortices, cingulate gyrus, as well as within subcortical structures, including the cerebellum (Gale *et al.*, 2005; Zappala *et al.*, 2012).

We also found that TBI patients incorporated the prefrontal cortex into the task-positive network, a network employed during performance of attention-demanding tasks (Fox *et al.*, 2005). This suggests more cognitive control and less automatic movement in TBI patients than in control participants.

**TBI-induced alterations to both structural and functional networks and association with behavior.** In many studies of brain networks, changes in functional or in structural network connectivity have often been associated with TBI, yet very few studies have addressed their combination. Here, we have shown that frontal brain areas in TBI patients demonstrate increased structural and functional connectivity as compared to control participants, and the activation at rest of the areas where connectivity increases is negatively correlated to postural control performance. This may refer to a compensatory *plasticity* mechanism that appears to suggest a different mode of balance control, namely increased controlled processing or less automatic processing of balance movements. Thus, it is apparently not a successful compensation whereby increased functional and structural connectivities leads to increased balance performance but rather a mandatory change in performance mode to be able to accomplish the balance tasks.

It is well known that frontal areas do not operate in isolation. In particular, it has been reported that interactions between the frontal cortex and the basal ganglia play a key role in movement control (Alexander *et al.*, 1990; Mink, 1996; Aron and Poldrack, 2006; Coxon *et al.*, 2010, Coxon *et al.* 2012; Hikosaka and Isoda, 2010). Thus, this fronto-striato-thalamic circuit enables frontal lobe regions to communicate with the basal ganglia, and is involved in a rich spectrum of different functions: motor and oculomotor circuits, executive functions, social behaviour and motivational states (Frank *et al.*, 2007). There is evidence that this reduced connectivity in the fronto-striato-thalamic circuit is correlated with a reduced subcortical grey matter volume and task performance after TBI (Leunissen *et al.*, 2014a, Leunissen *et al.* b).

Furthermore, it has been shown that white matter connectivity and subcortical gray matter volume continue to decrease up to 4 years post-injury (Farbota *et al.*, 2012), which might lead to a reorganization of the frontal brain regions in order to compensate for the damage to the fronto-striato-thalamic circuit. This potential response to the insult is in agreement with our findings (Leunissen *et al.*, 2014a, Leunissen *et al.* b). Previous studies have shown increased frontal connectivity to constitute a possible compensation mechanism after TBI. For example, using a working memory paradigm it was found that TBI patients required compensatory activation of the contralateral prefrontal region in order to perform certain tasks (Maruishi *et al.*, 2007). Resting state connectivity in frontal regions was also shown to increase in TBI patients (Palacios *et al.*, 2013), an increase that was correlated with cognitive performance.

## Summary

Using a new hierarchical atlas considering regions in relation to both structure and function, we demonstrated increased structural and functional connectivity in prefrontal regions in TBI patients relative to controls, relative to a general pattern of overall decreased connectivity across the TBI brain. Although this increased prefrontal connectivity reflected interactions between brain areas when participants are at rest, the enhanced connectivity was found to be negatively correlated with active behaviour (i.e., postural control performance). Thus, our findings obtained during rest do potentially reflect how TBI patients orchestrate task-related activations to support behaviour in everyday life. In particular, enhanced connectivity in TBI might help to overcome the deficits in cerebellar and subcortical connections, in addition to compensating for deficits when interacting with the task-positive network. Hence, it appears that there is greater cognitive control over certain actions in order to overcome deficits in their automatic processing.

## Acknowledgments

ID undertook a 2 month lab rotation to visit the laboratories of Swinnen and Marinazzo that was funded by the Health Department of the Basque Goverment. SS acknowledges financial support from Bizkaia Talent and European Commission through COFUND with the research project BRAhMS - Brain Aura Mathematical Simulation-(AYD-000-285). JMC acknowledges financial support from Ikerbasque: The Basque Foundation for Science and Euskampus at UPV/EHU. SPS was supported by FWO Vlaanderen (Levenslijn G.A114.11 and G.0708.14), the Research Fund KU Leuven (C16/15/070) and the Interuniversity Attraction Poles program of the Belgian federal government (Belspo, P7/11).

## Supplementary Material: Figure S1

**Supplementary Fig. 1:**
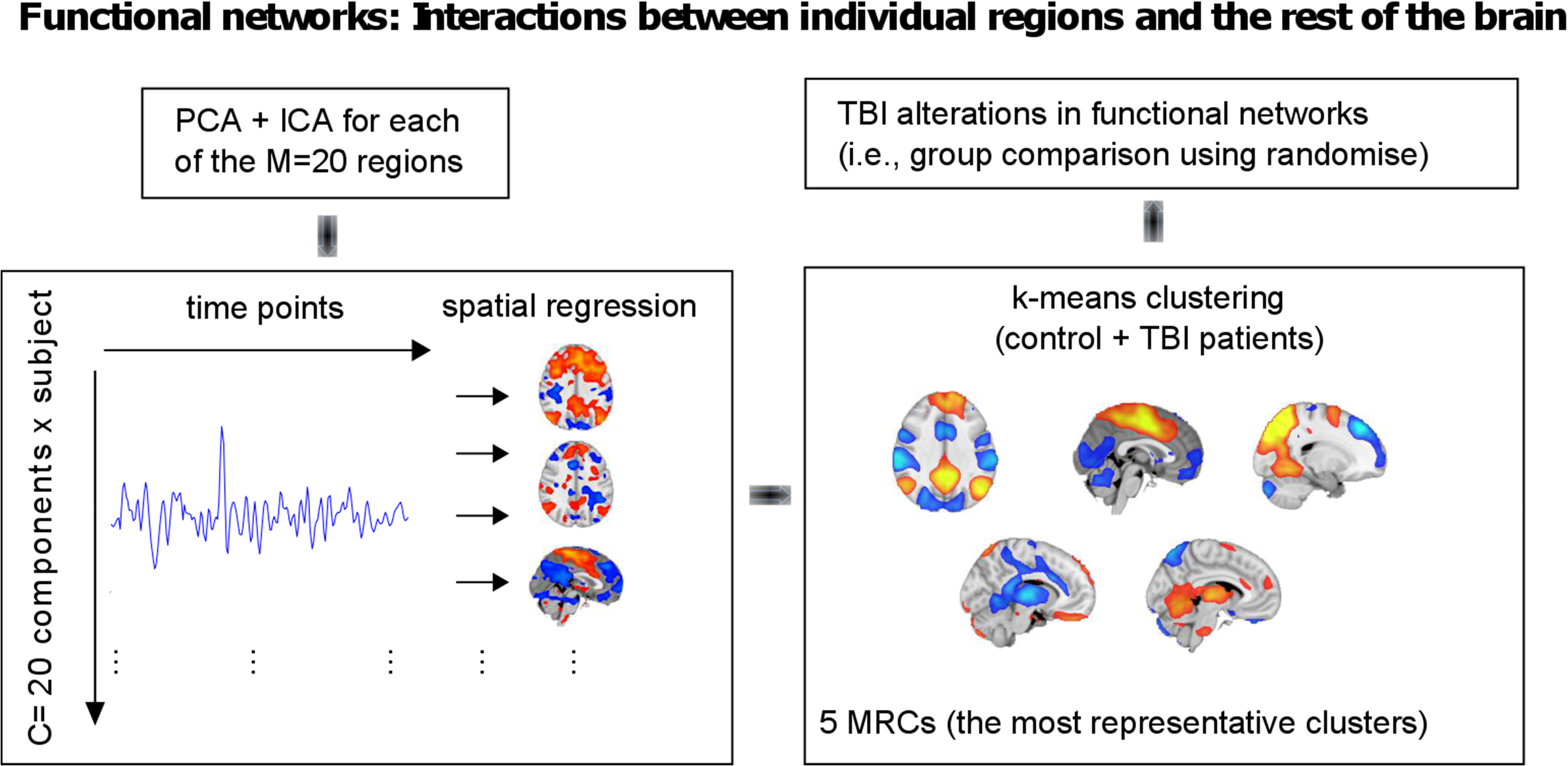
Sketch for the method to obtain functional interactions between individual regions and the rest of the brain. The alterations to the interaction network induced by TBI were addressed by applying a PCA+ICA to each of the M=20 regions in order to extract C=20 components for each region. These components were then spatially regressed to all the brains voxels in order to identify which voxels outside the region interacted most with each component (i.e.: to obtain the spatial map for each component). Finally, all the spatial maps were clustered using k-means, which provided the 5 MRCs (most representative clusters) that correspond to the 5 main networks that each of the M=20 regions interact with.

## References

Alexander G, Crutcher M, DeLong M (1990): Basal ganglia thalamocortical circuits: Parallel substrates for motor, oculomotor, “prefrontal” and “limbic” functions. Prog Brain Res 85:119146.

Amor TA, Russo R, Diez I, Bharath P, Zirovich M, Stramaglia S, Cortes JM, de Arcangelis L, Chialvo DR (2015): Extreme brain events: Higher-order statistics of brain resting activity and its relation with structural connectivity. EPL 111:68007.

Aron A, Poldrack R (2006): Cortical and subcortical contributions to stop signal response inhibition: role of the subthalamic nucleus. J Neurosci 26:2424e2433.

Barbey A, Belli A, Logan A, Rubin R, Zamroziewicz M, Operskalski J (2015): Network topology and dynamics in traumatic brain injury. Curr Opin Behav Sci 4:92–102.

Bishop C (2006): Pattern recognition and machine learning. Springer

Bonnelle V, Leech R, Kinnunen K, Ham T, Beckmann C, Boissezon XD, Greenwood RJ, Sharp DJ (2011): Default mode network connectivity predicts sustained attention deficits after traumatic brain injury. J Neurosci 31:13442–13451.

Bonnelle V, Ham TE, Leech R, Kinnunen KM, Mehta MA, Greenwood RJ, Sharp DJ (2012): Salience network integrity predicts default mode network function after traumatic brain injury. Proc Natl Acad Sci USA 109(12):4690–4695.

Caeyenberghs K, Leemans A, Heitger M, Leunissen I, Dhollander T, Sunaert S, Dupont P, Swinnen SP (2012): Graph analysis of functional brain networks for cognitive control of action in traumatic brain injury. Brain 135:1293–1307.

Caeyenberghs K, Leemans A, Leunissen I, Michiels K, Swinnen SP (2013): Topological correlations of structural and functional networks in patients with traumatic brain injury. Front Hum Neurosci 7:726.

Caeyenberghs K, Leemans A, Leunissen I, Gooijers J, Michiels K, Sunaert S, Swinnen SP (2014a): Altered structural networks and executive deficits in traumatic brain injury patients. Brain Struct Funct 219:193–209.

Caeyenberghs K, Siugzdaite R, Drijkoningen D, Marinazzo D, Swinnen SP (2014b): Functional Connectivity Density and Balance in Young Patients with Traumatic Axonal Injury. Brain Connect.

Craddock R, James G, Holtzheimer P, Hu X, Mayberg H (2012): A whole brain fmri atlas generated via spatially constrained spectral clustering, Hum Brain Mapp 33:1914–1928.

Craddock R, Jbabdi S, Yan C, Vogelstein J, Castellanos F, Martino AD, Kelly C, Heberlein K, Colcombe S, Milham MP (2013): Imaging human connectomes at the macroscale. Nat Methods 10:524–539.

Cordes D, Haughton V, Arfanakis K, Carew J, Turski P, Moritz C, Quigley MA, Meyerand ME (2012): Frequencies contributing to functional connectivity in the cerebral cortex in resting-state data. Am J Neuroradiol 22:1326–1333.

Coxon J, Goble D, Van I, De Vos J, Wenderoth N, Swinnen S (2010): Reduced basal ganglia function when elderly switch between coordinated movement patterns. Cereb Cortex 20:2368e2379.

Coxon J, Van Impe A, Wenderoth N, Swinnen S (2012): Aging and inhibitory control of action: cortico-subthalamic connection strength predicts stopping performance. J Neurosci 32: 8401e8412.

Damoiseaux J, Greicius M (2009): Greater than the sum of its parts: a review of studies combining structural connectivity and resting-state functional connectivity. Brain Struct Funct 213: 525–533.

Desikan R, Segonne F, Fischl B, Quinn B, Dickerson B, Blacker D, Buckner RL, Dale AM, Maguire RP, Hyman BT, Albert MS, Killiany RJ (2006): An automated labeling system for subdividing the human cerebral cortex on MRI scans into gyral based regions of interest. Neuroimage 31:968–980.

Diez I, Bonifazi P, Escudero I, Mateos B, Munoz M, Stramaglia S, Cortes JM (2015): A novel brain partition highlights the modular skeleton shared by structure and function. Scientific Reports 5:10532.

Drijkoningen D, Caeyenberghs K, Leunissen I, Linden C, Sunaert S, Duysens J, Swinnen SP (2015a): Training-induced improvements in postural control are accompanied by alterations in cerebellar white matter in brain injured patients. Neuroimage Clin 7:240–251.

Drijkoningen D, Leunissen I, Caeyenberghs K, Hoogkamer W, Sunaert S, Duysens J, Swinnen SP (2015b): Regional volumes in brain stem and cerebellum are associated with postural impairments in young brain-injured patients. Hum Brain Mapp 36(12): 4897–909.

Eickhoff S, Stephan K, Mohlberg H, Grefkes C, Fink G, Amunts K, Zilles K (2005): A new spm toolbox for combining probabilistic cytoarchitectonic maps and functional imaging data. Neuroimage 25:1325–1335.

Fagerholm E, Hellyer P, Scott G, Leech R, Sharp D (2015): Disconnection of network hubs and cognitive impairment after traumatic brain injury. Brain 138:1696–1709.

Farbota K, Sodhi A, Bendlin B, McLaren D, Xu G, Rowley H, Johnson SC (2012): Longitudinal volumetric changes following traumatic brain injury: a tensor-based morphometry study. Neuropsychol Soc 18–1006e1018.

Fox MD, Snyder AZ, Vincent JL, Corbetta M, Van Essen DC, Raichle ME (2005): The human brain is intrinsically organized into dynamic, anticorrelated functional networks. Proc Natl Acad Sci USA 102:9673–9678

Frank M, Scheres A, Sherman S (2007): Understanding decision-making deficits in neurological conditions: insights from models of natural action selection. Phil Trans R Soc B Biol Sci 362:1641e1654.

Gale S, Baxter L, Roundy N, Johnson S (2005): Traumatic brain injury and grey matter concentration: a preliminary voxel based morphometry study. J Neurol Neurosurg Psych 76:9848.

Gentry L, Godersky J, Thompson B (1988): Mr imaging of head trauma: review of the distribution and radiopathologic features of traumatic lesions. AJR Am J Roentgenol 150:663e672.

Godefroy O (2003): Frontal syndrome and disorders of executive functions, J Neurol 250:16.

Ham T, Sharp D (2012): How can investigation of network function inform rehabilitation after traumatic brain injury? Curr Opin Neurol 25: 662–669.

Hikosaka O, Isoda M (2010): Switching from automatic to controlled behavior: cortico-basal ganglia mechanisms. Trends Cogn Sci 14:154e161.

Hillary FG, Slocomb J, Hills EC, Fitzpatrick NM, Medaglia JD, Wang J, Good DC, Wylie GR (2011): Changes in resting connectivity during recovery from severe traumatic brain injury. Int J Psychophysiol 82(1): 115–123.

Hulkower M, Poliak D, Rosenbaum S, Zimmerman M, Lipton M (2013): A decade of dti in traumatic brain injury: 10 years and 100 articles later. AJR Am J Roentgenol 34:2064–74.

Kim J, Parker D, Whyte J, Hart T, Pluta J, Ingalhalikar M, Coslett HB, Verma R (2014): Disrupted structural connectome is associated with both psychometric and real-world neuropsychological impairment in diffuse traumatic brain injury. J Int Neuropsychol Soc 20(9):887–96.

Lancaster J, Woldorff M, Parsons L, Liotti M, Freitas C, Rainey L, Kochunov PV, Nickerson D, Mikiten SA, Fox PT (2000): Automated talairach atlas labels for functional brain mapping. Hum Brain Mapp 10:120–131.

Leunissen I, Coxon J, Caeyenberghs K, Michiels K, Sunaert S, Swinnen S (2014a): Subcortical volume analysis in traumatic brain injury: the importance of the fronto-striato-thalamic circuit in task switching. Cortex 51:67–81.

Leunissen I, Coxon J, Caeyenberghs K, Michiels K, Sunaert S, Swinnen S (2014b): Task switching in traumatic brain injury relates to cortico-subcortical integrity. Hum Brain Mapp 25:2459–69.

Levin H, Kraus M (1994): The frontal lobes and traumatic brain injury. J Neuropsychiatry Clin Neurosci 6:443–454.

Maki-Marttunen V, Diez I, Cortes J, Chialvo D, Villarreal M (2013): Disruption of transfer entropy and inter-hemispheric brain functional connectivity in patients with disorder of consciousness. Front Neuroinformatics 7:24.

Maruishi M, Miyatani M, Nakao T, Muranaka H (2007): Compensatory cortical activation during performance of an attention task by patients with diffuse axonal injury: a functional magnetic resonance imaging study. J Neurol Neurosurg Psychiatry 78(2): 168173.

Mayer AR, Mannell MV, Ling J, Gasparovic C, Yeo RA (2011): Functional connectivity in mild traumatic brain injury. Hum Brain Mapp 32(11): 1825–1835.

Miller E (2000): The prefrontal cortex and cognitive control, Nat Rev Neurosci 1:59–65.

Mink J (1996): The basal ganglia: focused selection and inhibition of competing motor programs. Prog Neurobiol 50:381e425.

Mori S, Crain B, Chacko V, van Zijl P (1999): Three-dimensional tracking of axonal projections in the brain by magnetic resonance imaging. Ann Neurol 45:265–269.

Newman ME (2004): Fast algorithm for detecting community structure in networks. Phys RevE 69:066133

Palacios E, Sala-Llonch R, Junque C, Roig T, Tormos J, Bargallo N, Vendrell P (2013): Resting-state functional magnetic resonance imaging activity and connectivity and cognitive outcome in traumatic brain injury. JAMA Neurol 70(7):845–851.

Park H, Friston K (2013): Structural and functional brain networks: from connections to cognition. Science 342:1238411.

Sharp D, Beckmann C, Greenwood R, Kinnunen K, Bonnelle V, Boissezon XD, Powell JH, Counsell SJ, Patel MC, Leech R (2011): Default mode network functional and structural connectivity after traumatic brain injury. Brain 134:2233–2247.

Sharp D, Scott G, Leech R (2014): Network dysfunction after traumatic brain injury. Nat Rev Neurol 10:156–166.

Smith DH, Meany DF (2000): Axonal Damage in Traumatic Brain Injury. Neuroscientist 6:1073–8584.

Smith S, Fox P, Miller K, Glahn D, Fox P, Mackay C, Filippini N, Watkins KE, Toro R, Laird AR, Beckmann CF (2009): Correspondence of the brains functional architecture during activation and rest. Proc Natl Acad Sci USA 106:13040–13045.

Sorensen T (1948): A method of establishing groups of equal amplitude in plant sociology based on similarity of species and its application to analyses of the vegetation on Danish commons. Kongelige Danske Videnskabernes Selskab 5:1–34.

Tagliazucchi E, Balenzuela P, Fraiman D, Chialvo D (2012): Criticality in large-scale brain fmri dynamics unveiled by a novel point process analysis. Front Physiol 3:15.

Tagliazucchi E, Carhart-Harris R, Leech R, Nutt D, Chialvo DR (2014): Enhanced repertoire of brain dynamical states during the psychedelic experience. Hum Brain Mapp 35:5442–5456

Tarapore PE, Findlay AM, Lahue SC, Lee H, Honma SM, Mizuiri D, Luks TL, Manley GT, Nagarajan SS, Mukherjee P (2013): Resting state magnetoencephalography functional connectivity in traumatic brain injury. J Neurosurg 118(6): 1306–1316.

Tzourio-Mazoyer N, Landeau B, Papathanassiou D, Crivello F, Etard O, Delcroix N, Mazoyer B, Joliot M (2002): Automated anatomical labeling of activations in SPM using a macroscopic anatomical parcellation of the MNI MRI single-subject brain. Neuroimage 15:273–289.

Vakhtin A, Calhoun V, Jung R, Prestopnik J, Taylor P, Ford C (2013): Changes in intrinsic functional brain networks following blast-induced mild traumatic brain injury. Brain Inj 27:1304–1310.

Van Impe A, Coxon JP, Goble DJ, Doumas M, Swinnen SP (2012): White matter fractional anisotropy predicts balance performance in older adults. Neurobiol Aging 33:1900–1912.

Wang R, Benner T, Sorensen A, Wedeen V (2013): Diffusion toolkit: a software package for diffusion imaging data processing and tractography. Proc Intl Soc Mag Reson Med 15:3720.

Zappala G, Schotten MTD, Eslinger P (2012): Traumatic brain injury and the frontal lobes: what can we gain with diffusion tensor imaging? Cortex 48:156e165.

Zhou Y, Milham MP, Lui YW, Miles L, Reaume J, Sodickson DK, Grossman RI, Ge Y (2012): Default-mode network disruption in mild traumatic brain injury. Radiology 265(3):882–892.

